# Photodynamic therapy effect on the ultrastructure of *Trichomonas vaginalis* trophozoites and their effectiveness in experimentally infected animals

**DOI:** 10.1101/327189

**Authors:** Thaisa H.S. Fonseca, Jessica M.S. Gomes, Marina Alacoque, Marcos A. Vannier-Santos, Maria A. Gomes, Haendel G.N.O. Busatti

## Abstract

**Background:** *Trichomonas vaginalis* is an amitochondrial parasitic that causes human trichomoniasis, the most common non-viral sexually transmitted infection in the world. The therapy of choice is metronidazole (MTZ). Despite MTZ effectiveness, resistant cases are becoming more frequent. Another point to emphasize are the side effects that may result in treatment discontinuation, leading to further spread of infection and emergence of resistant strains. This scenario reveals the need to develop new therapeutic options. Photodynamic therapy (PDT) is an experimental treatment that involves the activation of photosensitive substances and the generation of cytotoxic oxygen species and free radicals to promote the selective destruction of target tissues. A previous study, from our group, identified an excellent *in vitro* PDT activity using methylene blue and light emitting diode against MTZ sensitive and resistant strains of *T. vaginalis*. The aim of this study was to evaluate the efficacy of PDT *in vivo* and clarify its high trichomonicidal potential by evaluating its action upon *T. vaginalis* trophozoites through transmission electron microscopy (TEM).

**Methodology:** Seven-week-old female Balb/c mice were infected intravaginally with *T. vaginalis* trophozoites. On the third day of infection, methylene blue was introduced into the vaginal canal of the animals, which then received 68.1 J / cm^2^ of radiation for 35.6 sec. Control groups without infection and infected, treated with metronidazole were also included for comparison. Twenty-four hours after treatment the vaginal canal of the animals was scraped and the samples processed by the immunocytochemistry technique.
After in vitro photodynamic treatment, *T. vaginalis* trophozoites were processed for TEM. Ultrathin sections were collected in 400-mesh copper grids, contrasted with 5% uranyl acetate and 3% lead citrate, in aqueous solutions for 20 and 5 min., respectively and observed in a Jeol JEM 230 transmission electron microscope.

**Results:** TEM showed morphological changes such as centripetal displacement of organelles, cannibalism, hydrogenosomal damage, intense cytoplasmic vacuolization, dilated endoplasmic reticulum cisternae and membrane discontinuity, in both resistant and sensitive strains, suggesting that trichomonicidal activity is mainly due to necrosis.

PDT significantly reduced infection in animals treated with a single therapy session, compared to control groups, being statistically as efficient as MTZ.

**Conclusions:** Our results demonstrated high trichomonicidal activity of PDT with morphological alterations compatible with necrosis. Therefore these results indicate that PDT represents not only an alternative therapy for refractory trichomoniasis, but also routinely for this important neglected parasitic disease.

## Author summary

Trichomoniasis is the most prevalent non-viral Sexually Transmitted Infection (STI) worldwide. Although not causing death, it is associated with other potentially fatal STIs such as HIV. The success of their control is impaired by promiscuity, directing the efforts to restrain the transmission of disease to treatment. MTZ is the drug of choice. Despite its effectiveness, cases of resistance are frequent. It adds up to the resistance, the side effects that can interrupt the treatment culminating in more resistance and transmission of the disease. In this context, therapeutic alternatives may represent a solution not only to solve the increasing resistance, but also for trichomoniasis control. We evaluated the efficacy of photodynamic therapy (PDT) as trichomonicide in experimentally infected animals and its action on the parasite was evaluated by transmission electron microscopy (TEM). We use methylene blue as a photosensitizing agent and as light source, a diode-emitting red light. PDT showed expressive trichomonicidal activity in vivo, absence of toxicity and excellent applicability, both in sensitive and resistant strains of *T vaginalis*. TEM revealed morphological alterations indicative of cell death by necrosis. Therefore, PDT proved to be a feasible therapeutic alternative, especially with regard to its achievability. In addition, compared to other therapies, PDT has the advantage of having a high selectivity, as the light sources are directed to the lesion site; has low possibility of developing photo-resistant strains; its trichomonicide action is fast; has a low risk of induction of mutagenic effects and still has low cost.

## Introduction

*Trichomonas vaginalis*, the causative agent of human trichomoniasis, is not only the most prevalent non-viral sexually transmitted pathogen in the world [1], but also enhances human immunodeficiency virus (HIV) transmission and it can also lead to cervical and prostate cancer [2].

Nitroimidazoles, such as metronidazole (MTZ), have been the drugs of choice for the treatment of the disease [3]. However, the success of the treatment is limited by the side effects that lead to the discontinuation of the therapeutic regimen [4], promoting the emergence of resistant isolates, impairing the control of the disease [5]. This scenario indicates the great need for the development of new anti-*Trichomonas* therapeutic options for a more efficient control of the disease, reduction of the risks to the patients health, besides the reduction of the emergence of resistant strains.

PDT is an experimental treatment with great therapeutic potential for neoplastic and non-neoplastic diseases [6]. As a basic principle of the technique, visible light activates a photosensitizing substance accumulated in tissue or organisms [7]. The interaction between the excited photosensitizer in the presence of molecular oxygen triggers the production of singlet oxygen (^1^O_2_) as well as other reactive oxygen species (ROS) [8], inducing cell death by necrosis or apoptosis [9]. Several cellular organelles/components can comprise targets, such as nucleic acids, plasma membrane, mitochondria, endoplasmic reticulum and cytoskeletal structure [10].

In a previous study, our group demonstrated the excellent trichomonacidal activity of PDT using methylene blue [11]. High reduction of trophozoite numbers (approximately 90%) was observed with only one *in vitro* photodynamic treatment session. High anti-*T. vaginalis* activity was observed upon a MTZ-resistant isolate, point out the use of PDT as an alternative therapy in trichomoniasis.

Thus, here we evaluated the efficacy of PDT in experimentally infected animals and approached its mechanism of action on *T. vaginalis* cells.

## Methods

### T. vaginalis

A strain isolated from a 40-year-old symptomatic female (Rio de Janeiro, Brazil) and CDC 085 strain (ATCC 50143), isolated in Columbus, Ohio, from a treatment-refractory patient [3]. Both were maintained in YI-S medium [12] at 37 °C under axenic cultivation.

### Photosensitizer

The photosensitizer employed was methylene blue (Sigma, St. Louis, MO, USA) dissolved in sterile Mili-Q water at a stock concentration of 0.25% and stored in the dark at 4 °C until use.

### *In vitro* treatment with MB and PDT

Initially, a suspension of 4 × 10^5^ trophozoites from each strain was individually transferred to plastic microtubes at a final volume of 1 mL to enable treatment. The Light Control (LC), Methylene Blue (MB), and Photodynamic Therapy (PDT) groups were compared to the Control (CTR) group. Increasing concentrations of the photosensitizer (25 μM, 100 μM, and 250 μM) were added to the trophozoites of the MB and PDT groups for 30 min (pre-irradiation time) in the dark, in a bacteriological incubator at 37°C. Next, the mixture was washed by centrifugation (850 × *g*, 10 min, 4°C) with 0.5 mL phosphate buffered saline at pH 7.2 (PBS - Sigma-Aldrich^®^, USA) to remove unabsorbed photosensitizer excess. The contents were then transferred to 13 × 100 mm glass tubes (Pyrex^®^, Corning, Inc., Corning, NY, USA). Individually, each LC and PDT tube received irradiation from the bottom up at three energy density range (E.D: 68.1 J/cm^2^; 95.45 J/cm^2^ and 122.7 J/cm^2^). At the end of this process, YI-S medium was added to all tubes to a final volume of 6 mL. The tubes were incubated for 24 h in a bacteriological incubator at 37°C. After incubation, the viability and quantification of the trophozoites were determined using a flow cytometer (FACScan, Becton Dickinson, Franklin Lakes, NJ, USA) with a 1 mg/mL concentration of propidium iodide (PI - Thermo Fisher Scientific, USA). The analysis was performed using Cell Quest ProTM software (Becton Dickinson, USA), and 50.000 events were acquired for each sample inside the gate. All experiments were performed in triplicate and repeated at least two times.

### Transmission electron microscopy (TEM)

After *in vitro* photodynamic treatment, *T. vaginalis* trophozoites were fixed for 1 hour at room temperature in 2.5% glutaraldehyde solution in 0.1M sodium cacodylate buffer, pH 7.4. After fixation, the parasites were washed, postfixed in 1% osmium tetroxide and 0.8% potassium ferrocyanide and 5 mM calcium chloride in the same buffer at room temperature, in the dark. The cells were then washed in this buffer, dehydrated in increasing concentrations of acetone and embedded in PolyBed resin (Polyscience, Warrington PA). Ultrathin sections obtained with a diamond knife were collected in 400-mesh copper grids, contrasted with 5% uranyl acetate and 3% lead citrate, in aqueous solutions for 20 and 5 min., respectively and observed in a Jeol JEM 230 transmission electron microscope.

### *In vivo* photodynamic therapy

For the irradiation stage of the animal’s vaginal canals, we adapted a removable fiber-optic tip 2mm in diameter to the LED device, assuring suitable and efficient local irradiation. The final power of the apparatus was 60 mW (λ: 630 nm; E.D: 68.1 J/cm^2^; t: 35.6s). Seven-week-old female Balb/c mice were divided into the following experimental groups, containing 8 animals each: CTR (trophozoites only), MTZ CTR (trophozoites + MTZ treatment), LC (trophozoites + LED light 68.1 J/cm^2^), MB (trophozoites + 100μM MB) and PDT (trophozoites + 100μM MB + 68.1 J/cm^2^ LED light). The animals were kept at 22–24 °C and 70% humidity with free access to water and food under a light / dark cycle of 12:12 h.

The pre-estrogenization procedure followed the protocol standardized by [13]. Then, on day 0 and day 1 all animals were infected intravaginally with 1 x 10^7^ *T. vaginalis* trophozoites, strain JT (sensitive to MTZ). In a previous study [15], this strain was shown to be significantly more resistant to PDT than the CDC 085 strain (resistant to MTZ), so we employed it for the *in vivo* assays.

On day 2 post-infection (day 3) a solution of 100 μM MB was inoculated into the vaginal canal of the MB and PDT groups with the aid of a sterile plastic tip micropipette. Animals of the CTR and LC groups received sterile PBS intravaginally. Ten minutes after substance association (pre-irradiation time), animals from PDT and LC groups were irradiated (68.1 J/cm^2^) for 35.6 sec. The MTZ CTR group received treatment via gavage with MTZ single dose (2g/kg) on the second post-infection day.

24h after treatments, the vaginal samples were collected by scraping and processed for the immunocytochemistry technique, as previously described [14]. For each experimental group, the trophozoites were quantified in ten random microscopic fields the intensity of the infection was considered intense (above 12 trophozoites per field), moderate (5 to 11 trophozoites per field) and low (1 to 4 trophozoites per field).

### Ethics Statement

All protocols conducted with animals were designed and carried out in accordance with international ethical standards for animal experimentation (Helsinki Declaration and its amendments) and were approved by the Committee on Ethics in Animal Experimentation of the Federal University of Minas Gerais (Protocol number 140/2017). All efforts were made to minimize animal suffering and to reduce the number of animals used.

### Statistical analysis

The results obtained in the *in vivo* photodynamic therapy assays were analyzed by the non-parametric Kruskal-Wallis test and the Dunn post-test (p < 0.05). The analysis were performed in software Prism 6.0 (GraphPad, San Diego CA).

## Results

Microscopic analysis showed that the parasites were blue MB-stained, indicating that the 30 minutes exposure was sufficient for the photosensitizer to react with the protozoan compartments (Fig 1).

**Fig 1.**
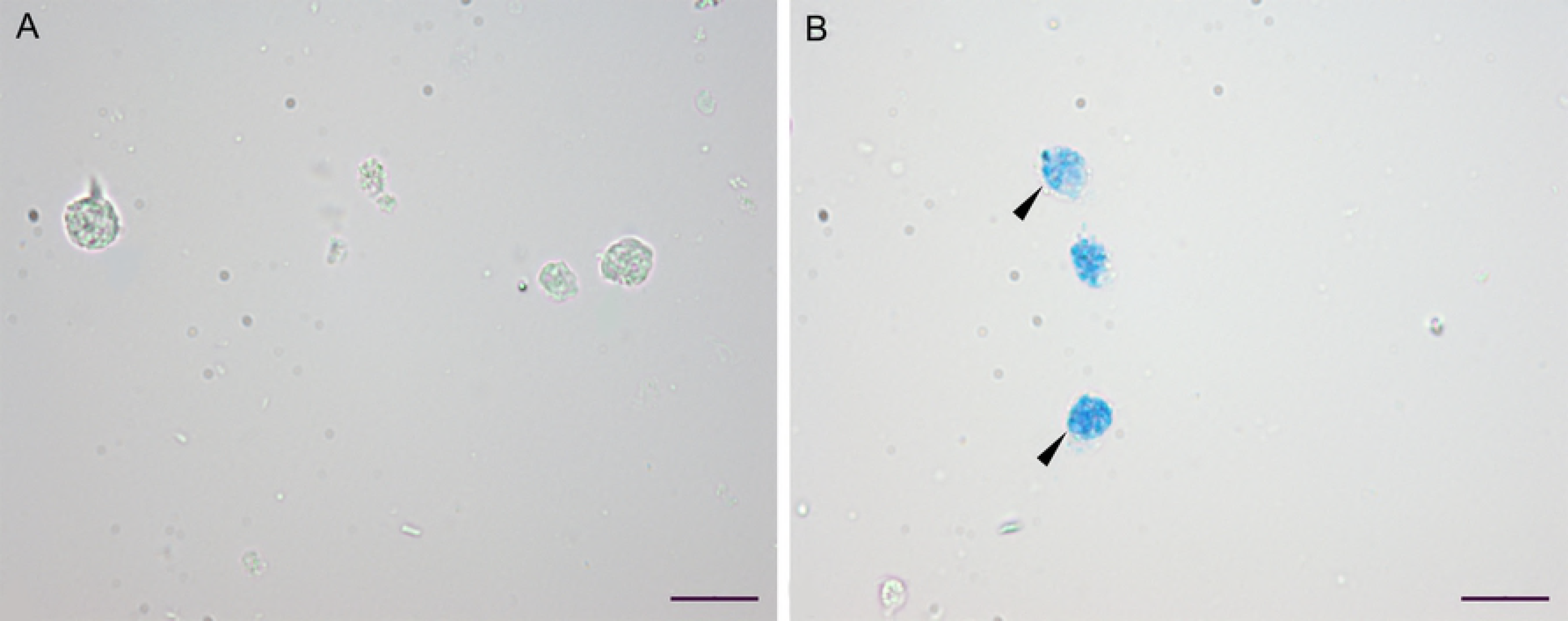
Marking of *T. vaginalis* trophozoites. Unlabelled (A) and 250 μM methylene blue treated for 30 minutes in the dark (B) *T. vaginalis* trophozoites (arrowheads). Bar: 50μM.

### Ultrastructural changes in trophozoites of *T. vaginalis*

In order to determine the effects caused by MB and PDT on *T. vaginalis* trophozoites, *in vitro* treatment kinetics were performed using distinct concentrations of MB in the absence and presence of irradiation (68.1 J/cm^2^). TEM was employed to evaluate the treatments-induced damages on the sensitive and resistant strains after 12h and 24h of incubation.

### Sensitive strain

The *T. vaginalis* cells of the untreated control (CTR) and light control (LC) groups presented well-preserved subcellular structures and cytoplasmic organelles, such as typical and numerous hydrogenosomes, endoplasmic reticulum cisternae, costa, nuclei, homogeneous cytoplasm with the presence of vacuoles, Golgi apparatus and axostyle (Fig 2A and 2B).

**Fig 2.**
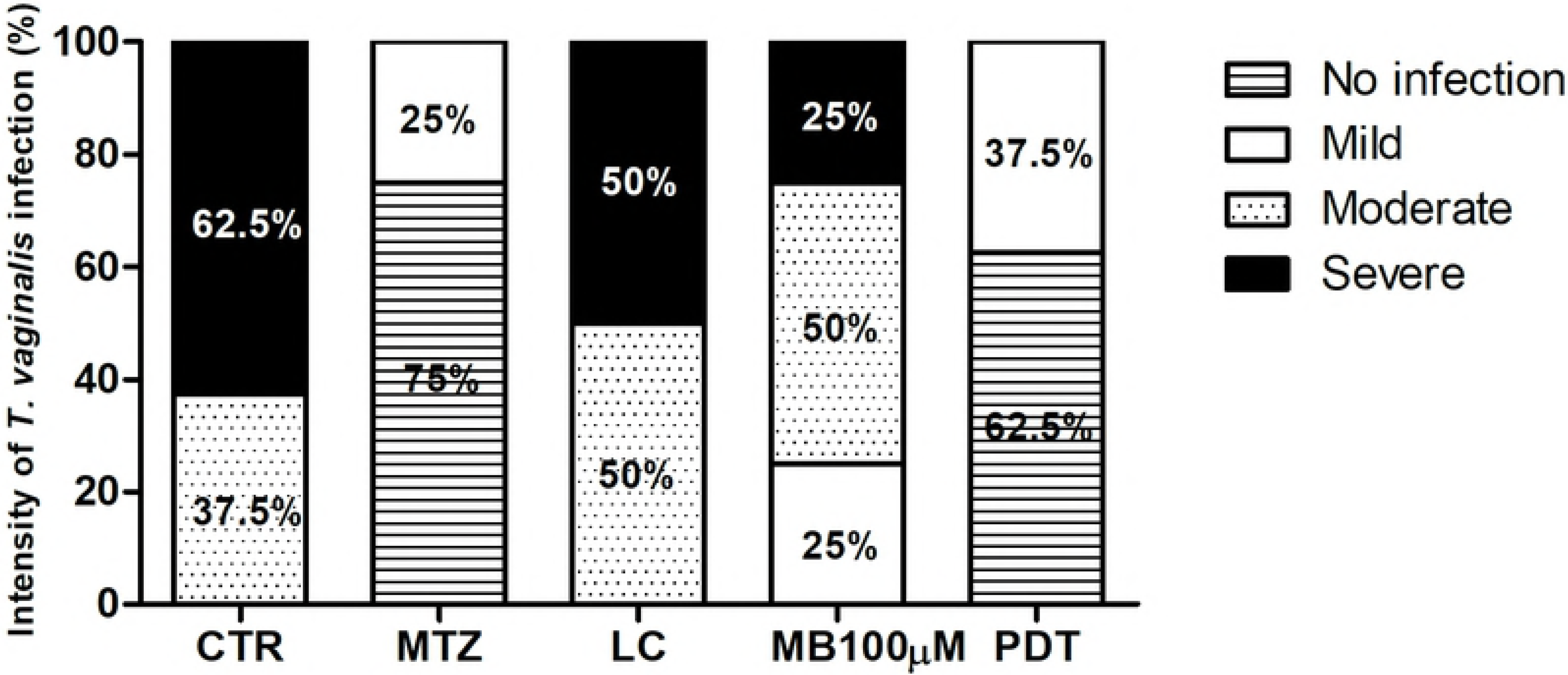
Changes observed in trophozoites of the JT strain after treatment with MB and PDT. Unlike control (A) and light control (B) cells that had well-preserved cytoplasm, some parasites treated with MB alone demonstrated centripetal displacement of organelles (C), numerous glycogen deposits (arrowheads) and moderate vacuolization, eventually displaying membranous material (D, arrows). In the PDT groups, it was possible to observe cannibalism by the uptake of hydrogensomes (E, arrowheads), vacuolization with granular material (E, arrow), hydrogenosome uptake (E, arrowheads) and, large autophagic vacuoles with large multilayered membrane-bounded compartments (F, arrows) and dilated endoplasmic reticulum cisternae (F, arrowheads). Many cells presented multivesicular bodies in the cell periphery (G, arrow, lower inset) that eventually fused with the parasite plasma membrane (G, upper inset) and PDT effects culminated in structural disorganization and remarkable reduction of cytoplasmic electrondensity (H).

In the presence of only MB, the trophozoites exhibited several changes in their ultrastructure. Some parasites displayed centripetal relocation of the organelles to the inner (more anaerobic) portions of the cytoplasm (Fig 2C), accumulation of numerous glycogen granule aggregates, dilated hydrogenosomal peripheral vesicles, moderate vacuolation and slightly reduced cytoplasmic electron density (Fig 2D).

In the PDT groups, process of phagocytosis of hydrogenosomes from the extracellular medium (Fig 2E) associated with a cytoplasmic exclusion zone characterizing the cytoskeleton participation in the parasite cannibalism. In addition, the presence of large and numerous autophagosomes with high cytoplasm-derived content, highly extracted cytoplasmic matrix, small expansion in the Golgi *cis* region with many budding vesicles, low hydrogenosome numbers, and dilated endoplasmic reticulum cisternae are prominent (Fig 2F).

Many trophozoites presented membrane-containing autophagosomes located at the periphery of the cells and fusing with the plasma membrane (Fig 2G). The advanced PDT effects included intense cellular damage, with structural disorganization and intense vacuolization, membrane discontinuity, extensive cytoplasmic extraction, and large debris assemblies (Fig 2J).

### Resistant strain

The resistant trophozoites CTR and LC groups, presented typical hydrogenosomes, but with reduced dimensions as compared to the sensitive strain, normal cytoskeleton and intact organelles including nuclei, which were eventually multiple (Fig 3A and 3B).

**Fig 3.**
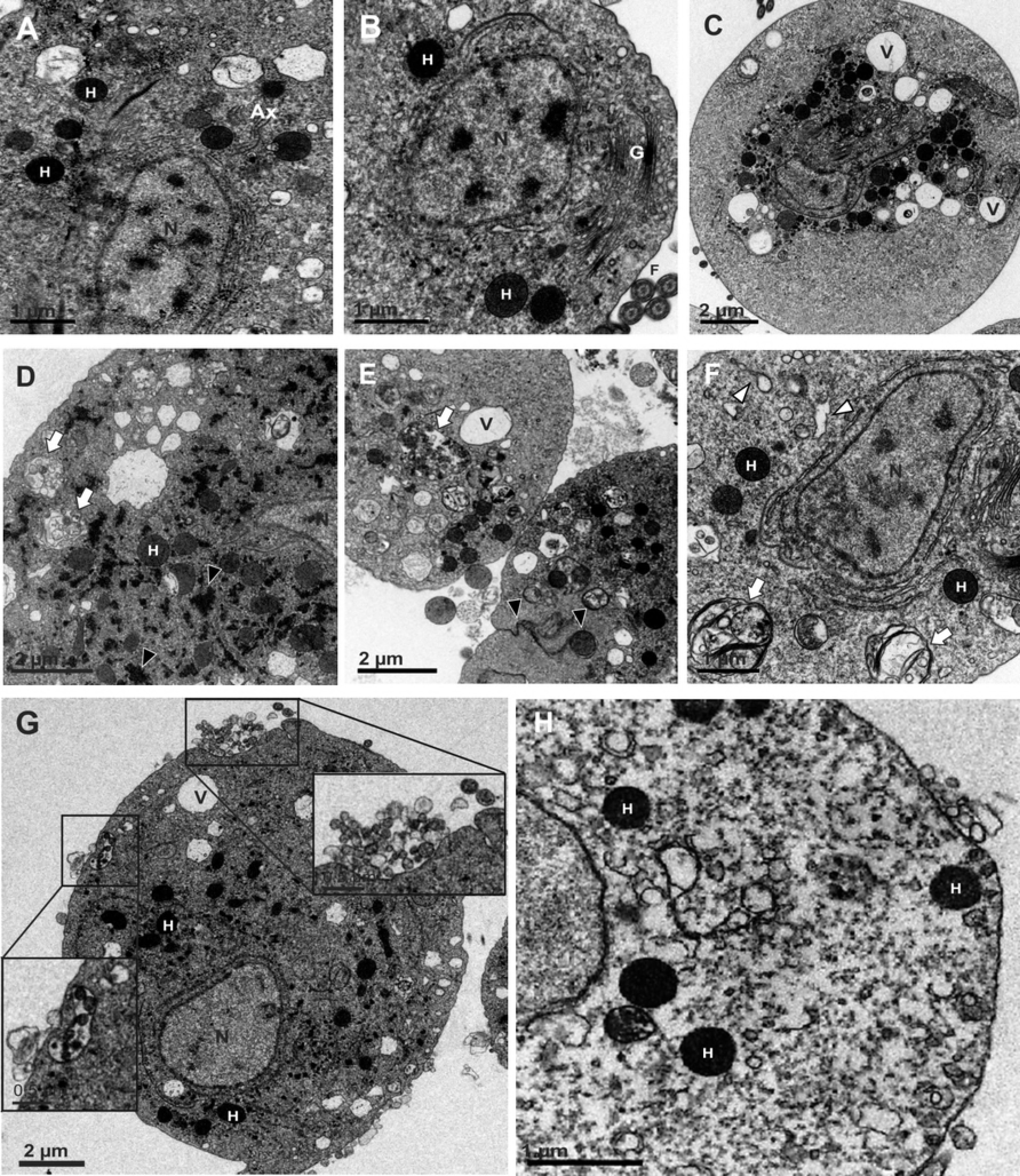
Trophozoites of CDC 085 strain after treatment with MB and PDT. Control cells (A) and light control (B) showed well-preserved hydrogensomes (H), digestive vacuoles (black arrowheads) and dilated endoplasmic reticulum cisternae (white arrowheads), as well as multinucleated (N) trophozoites (B). Cells incubated with MB (C) and submitted to PDT (D) exhibited hydrogensomal damage. Photodynamic treatment also led to the formation of autophagosomes, characterized by multiple membranes involving cytoplasmic portion (E), as well as events of cannibalism among parasites (F, black arrows; G, white arrow). Parasites submitted to PDT also demonstrate exocytosis of granular material (G, thick black arrows), engulfment (thin black arrows) and internalization (thick white arrow) by sibling parasite. Cell surface interdigitation was observed (G, thin white arrow). Trophozoite differentiation in pseudocysts was indicated by flagellar (H, arrow) internalization.

The presence of MB, without irradiation, led to decreased hydrogenosomal electrodensity and dilated endoplasmic reticulum cisternae (Fig 3C).

The early effects of PDT are marked by granular hydrogenosome matrix and high vacuolization, followed by membrane discontinuity leading to cytoplasmic extraction (Fig 3D). We also observed the formation of large myelin-like Figs containing cytoplasmic material with glycogen granules (Fig 3E).

A frequently observed event in the resistant strain, apparently exacerbated by PDT, was the formation of numerous surface interdigitations among adjacent trophozoites, indicating extensive cannibalism associated with necrotic areas. Both exocytosis and endocytosis of granular material were observed (Fig 3G), and internalized flagella were also detected (Fig 3H).

### PDT activity *in vivo*

The effects of photodynamic therapy *in vivo* treatment were determined by the immunocytochemical analysis of vaginal scrapings of animals experimentally infected with *T. vaginalis.* We did not observe a significant difference between the CTR or LC and MB groups. Nevertheless, the MTZ and PDT groups presented significantly (p<0.05) lowered mean parasite load, as compared to controls.

The intensity of the infection determination revealed that 62.5% of the animals in the CTR group had intense infection and 37.5% of the moderate type. Treatment with a single dose of MTZ resulted in healing of 75% of the animals, with the remaining 25% being mildly infected. Half of the animals in the LC group had intense infection whereas the other half had a moderate one (Fig 4). In addition, 25% of the animals in the MB group had intense, 50% moderate and 25% mild type infections. The animals in the PDT group showed 37.5% mild infection and 62.5% were absent. These data demonstrate that PDT significantly (p < 0.05) reduced infection in animals treated with a single treatment session, being statistically as efficient as the MTZ group (Fig 4).

**Fig 4.**
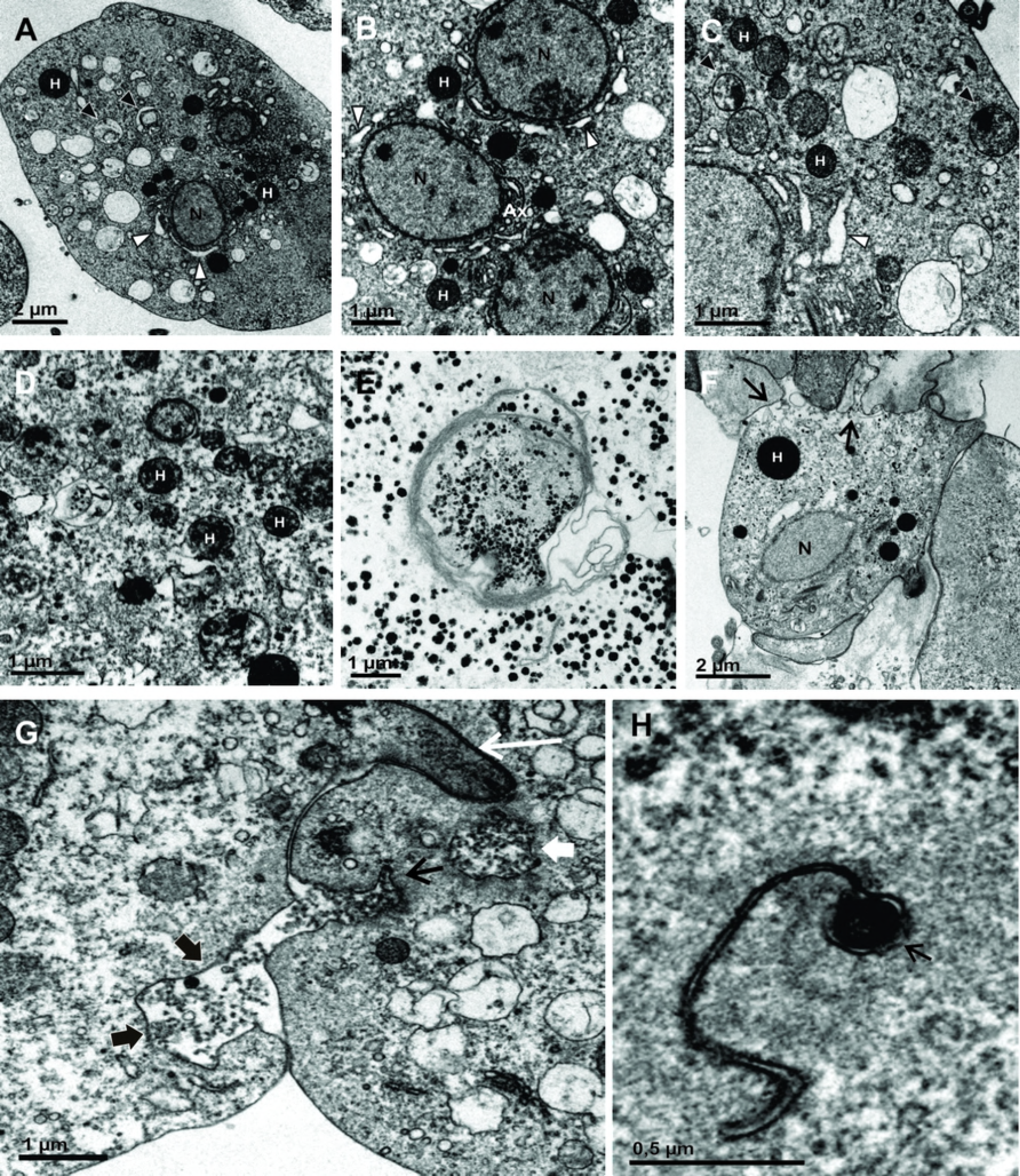
Intensity of infection by *T. vaginalis in vivo* treatments. The effects of photodynamic treatment *in vivo* were determined by the immunocytochemical analysis of vaginal scrapings in experimentally infected animals, according to the following groups: CTR (trophozoites only), MTZ (trophozoites + MTZ treatment), LC (trophozoites + LED), MB (trophozoites + 100μM MB) and PDT (trophozoites + 100μM MB + LED). The classification of the infection was standardized according to the profile: No infection = no trophozoite / field; Mild = 1 to 4 trophozoites / field; Moderate = 5 to 11 trophozoites / field; Severe = 12 or more trophozoites / field.

## Discussion

Nitroimidazoles such as metronidazole (MTZ) is a widely used antimicrobial agent against several infections and have been the therapeutic choice for the treatment of trichomoniasis [3]. Despite being one of the most prescribed medications, patients treated with MTZ have presented several undesirable side effects that can culminate in the discontinuation of the therapeutic scheme, promoting the rise of resistant strains [4]. Thus, therapeutic alternatives, including analogues [15], new agents [16] and regimens such as PDT [11] are desirable.

Compared to other therapies, PDT has the advantage of having great selectivity, since affected cells or tissues may comprise specific targets as light sources are directed to the lesion site [17]. Moreover, such therapy is also characterized by the low possibility of developing photoresistant strains, fast acting and low risk of induction of mutagenic effects [6].

In previous work, our group confirmed the efficacy and safety of photodynamic treatment with MB on susceptible and resistant strains of *T. vaginalis* [11]. Among the photosensitizers currently used, MB, a phenothiazine compound, has proven PDT-associated activity [18,19]. In this work, we used LED technology that, although less common in PDT, proved to be a good alternative in the treatment of cutaneous and mucosal wounds [20].

However, this is the first report of the effects caused by MB and PDT on the ultrastructure of *T. vaginalis* trophozoites and their effectiveness *in vivo*. Here we evaluated the damage caused to MTZ sensitive and resistant trophozoites by TEM.

In the sensitive strain, the trophozoites of the CTR and LC groups showed usual ultrastructure, presenting intact cytoplasmic organelles. However, in the presence of MB, without irradiation, we observed a centripetal migration of the organelles to the more anaerobic inner portion of the cells. It is possible that oxidative stress functions as a stimulus, causing the organelles such as hydrogenosomes to associate to the axostyle, and leaving large regions of organelle-free ectoplasm. Similar effect was observed in *Giardia lamblia* [15] trophozoites incubated with MTZ demonstrated evident ultrastructural disorganization characterized by the centripetal displacement of peripheral vesicles and internalization of the cytoskeletal components of the flagella and adhesive disc in the cytoplasmic matrix.

MB also caused the accumulation of numerous masses of glycogen throughout the cytoplasm. These glycogen masses may have occurred in response to the stress generated by PDT, since glycogen production and accumulation can reach up to three times the basal level in response to thermal shock, hyperosmotic and oxidative stress [21]. In addition, stress-induced glycogen accumulation, and limited fermentation capacity of *Saccharomyces cerevisiae*, results in the yeast adaptive mechanisms such as acquisition and accumulation of the reserve carbohydrates such as glycogen and trehalose [22].

Also approaching the sensitive strain, we observed that PDT caused considerable dilatation of endoplasmic reticulum (ER) cisternae, indicating that the oxidative stress caused by PDT may have interfered in the calcium homeostasis and/or cytochrome P450 activity of these parasites. Some drugs affect the cytoskeleton and can modify the ER, the Golgi apparatus and the distribution of vesicles in most cells. The drug-altered *Leishmania* ER can be found both in contact with the plasma membranes of the parasite and in cell surface protrusions [23].

Surviving cells can feed on the nutrients released by dead sibling cells or on attacking the cells *per se*, in a behavior termed “cannibalism” [24]. Microbial cannibalism was reported in cilliated protozoa [25], dinoflagellates [26] and *T. vaginalis* [27], characterized by the attack of living cells of the same strain. In the PDT groups, such mechanism of phagocytosis of nutrients from the external environment may comprise an energy source for parasite survival.

In addition, parasites of the sensitive strain exposed to PDT presented intense autophagy, numerous vesicles, plasma membrane discontinuity and reduced cytoplasmic electrondensity, associated with large debris amounts were also observed after PDT. The *in vitro* activity of PDT presented greater inhibitory capacity as compared to MB group (p < 0.05) [11].

Autophagy may comprise both processes for cellular homeostasis and cell death [28]. ROS can cause irreversible oxidative damage to proteins, lipids and DNA representing the main source of damage in biological systems. Autophagy plays an important role in the recycling of these biomolecules and can be considered an effective repair/escape system. Thus, low ROS concentrations play a role as signaling molecules triggering autophagy [29].

Parasites from resistant strain, CTR and LC groups, presented typical hydrogenosomes, but less numerous than the sensitive strain, normal cytoskeleton elements (*e.g.* costa and axostyle), small vacuoles and normal Golgi. However, in these groups the presence of dilated ER cisternae and multinucleation were observed. *Vinca* alkaloids produced a reversible block of cytokinesis in *Trypanosoma cruzi* with the presence of multiple nuclei and kinetoplasts [30]. Multinucleated and multiflagellated cells indicate a truncated or hampered cytokinesis in vinblastine-resistant *Leishmania amazonensis* [31]. Interestingly, the multinuclear phenotype may promote drug resistance in malignant cells [32]. Thus, *T. vaginalis* multinucleation may be implicated in metronidazole-resistance by the protozoan.

In the presence of MB, non-irradiated parasites of the resistant strain, presented aggregated electron-dense material, loss of cytoplasmic matrix and significantly dilated ER cisternae. Inactive hydrogenosomes can be readily recognized by matrix electron-dense deposits, termed nucleoids [33].

Hydrogenosomes are organelles involved in catabolic processes that include glycolysis. The oxidative decarboxylation of pyruvate, linked to ferredoxin-mediated electron transport is involved in ATP generation and induces the activation of the 5-nitroimidazoles, including MTZ [34]. This redox organelle may be destroyed by treatment of *T. foetus* with the putrescine analogue 1,4-diamino-2-butanone [35].

The hydrogenosomes usually present a smaller size in resistant strains when compared to the sensitive strains, being able to be approximately 20% smaller [36]. In addition, the typical electron density of *T. vaginalis* hydrogenosomes, largely due to the Fe-S pool, is not evident in resistant cells. This fact is in agreement with the negative regulation of the hydrogenosomal function [34, 36].

As observed in the sensitive strain, MB groups did not demonstrate the same extent and severity of damage as compared to PDT groups in the resistant strain.

Among the PDT groups of resistant strain, events of extensive cannibalism were frequently observed, characterized by numerous interdigitations among adjacent trophozoites associated to areas of necrosis. Such behavior may comprise an intraspecific competition mechanism triggered by starvation stress [25]. In addition, this mechanism could control the parasite load, possibly leading to asymptomatic infections [27].

Alterations in ER of trypanosomatids may lead to the formation of myelin-like bodies, which are often associated with oxidative stress and autophagy [23]. Similar changes were observed following PDT on resistant strain. It is noteworthy that this structure displayed cytoplasmic content including glycogen granules, presumably comprising autophagosomes. These membranes may be derived from ER, nucleoplasmic reticulum [37] or mitochondrial membranes [38].

*T. foetus* submitted to stress conditions internalize their flagella to form pseudocysts, and may be an escape strategy from environmental changes [39]. This event may be a temporary form, representing an early stress-induced event, prior to cell death. The formation of pseudocysts was also observed in this study after PDT on resistant strain.

PDT mediated death is influenced by the cell type, identity and concentration of the photosensitizer and light-doses used in the PDT protocol [40]. Treatment with high energy density and high concentrations of the photosensitizing substance leads to cell death by necrosis, whereas low dose treatment tends to induce apoptosis. Depending on the amount of ROS produced, death by autophagy may also occur [41].

The changes caused by PDT in the parasite *T. vaginalis* are different from those caused by MTZ, already described in the literature. Nielsen [42] did not observe a difference in the hydrogenosomes, called at the time of “chromatic granules”, describing only digestive vacuoles that may in part correspond to the autophagic vacuoles. Our work has shown, in addition to the autophagic vacuoles, the destruction of the hydrogenosomes that can result from oxidative stress. In view of the ultrastructural alterations described here, we observed evidence of necrotic cell death caused by PDT, on both sensitive and resistant *T. vaginalis*, in a manner analogous to that reported previously [43].

Furthermore, the efficacy of *in vivo* PDT was demonstrated in this study, with the parasite load in the PDT group being significantly reduced with only 1 treatment session (37.5% of the animals had mild infection and 62.5% absent), thus indicating that PDT may comprise not only as alternative therapy for refractory cases, but also as therapy of choice for this neglected parasitic disease.

## Acknowledgments

We thank Dr. Fernando Costa e Silva Filho for providing the JT strain of *T. vaginalis*.

